# A resurrection ecology test for the effects of ionizing radiation and other environmental correlates on *Daphnia* evolution

**DOI:** 10.1101/2023.02.10.524566

**Authors:** Twyla Neely-Streit, Megan C. O’Toole, Emily M. Ciolak, Musharrat Islam, Adam J. Heathcote, Andrew L. Masterson, Matthew R. Walsh, Yoel E. Stuart

**Affiliations:** Department of Biology, Loyola University Chicago, Chicago, IL, USA; St. Croix Watershed Research Station, Science Museum of Minnesota, Marine on St. Croix, MN, USA; Stable Isotope Biogeochemistry Laboratory, Northwestern University, Evanston, IL, USA; Department of Biology, University of Texas Arlington, Arlington, TX, USA

**Keywords:** Cesium-137, Ephippia, Lake, Lead-210, Limnology, Nevada Test Site, Nuclear Fallout, Paleolimnology, Radioisotope Dating, Sediment Core, Stable Isotopes, Utah

## Abstract

Acute exposure to ionizing radiation has well-documented, immediate negative consequences for individuals. However, the evolutionary consequences for populations exposed to ionizing radiation is unclear. For example, a meta-analysis of taxa exposed to Chernobyl fallout found some evidence for elevated mutation rates in animal and plant taxa; however, in people, *de novo* mutation rates in offspring of parents exposed to radiation during and after the Chernobyl accident were no higher than controls. Furthermore, whether irradiation and increased mutation rates drive adaptation to radiation also has mixed support. Ambiguity in both cases likely arises from the difficulty of studying mutation rates and adaptation after rare nuclear events whose ionizing radiation is distributed heterogeneously in time and space. Here, we report an attempt to better address this difficulty with a “resurrection ecology” study of *Daphnia spp*. in Utah lakes that experienced nuclear fallout from US Department of Energy weapons testing in the 1950s and 1960s. The idea was to recover dormant *Daphnia* eggs from sediment cores that spanned the nuclear testing era in the American West. We predicted that survival and fecundity of eggs hatched in the lab would show fitness declines correlated with ionizing radiation fallout and a potential recovery once nuclear testing stopped. We successfully obtained multiple cores from three lakes that dated back to the 1800s. We isolated >4700 dormant eggs from those cores, spanning the nuclear era, but were only able to hatch a single egg in the lab. Thus, we could not conduct life history experiments to test our prediction. The purpose of this manuscript, therefore, is to describe the study and make our radioisotope core dating and sedimentation data available to other paleolimnological researchers. We also report a side study of stable isotope change through time measured from dormant eggs and the sediment.

## Introduction

One proposed start date for the Anthropocene is the 1945 detonation of the first atomic bomb, which introduced a qualitatively new way for humans to alter the Earth. The radiation spike from worldwide, above-ground weapons testing in the mid-1950 and 1960s will likely be a reliable marker of human presence for hundreds of thousands of years. Whether human-induced radioactivity will leave as clear an imprint on biological evolution is an open question.

Human-caused irradiation could influence evolution in at least two ways (Polikarpov 1998). First, it might increase mutation rates, though the evidence for this phenomenon is mixed. For example, Yeager et al. (2021) found no relationship between people exposed to ionizing radiation at Chernobyl and the rate of *de novo* mutation in their offspring. On the other hand, a meta-analysis of populations of plants, animals, and microbes from the Chernobyl Exclusion Zone (CEZ) found a positive correlation between radiation exposure and mutation rate (r = 0.67, 95% confidence interval 0.59-0.73; 151 effect sizes spanning 45 studies and 30 species; Møller and Mousseau 2015).

Second, to the extent that increased mutation rates introduce additional deleterious and beneficial variants, they should affect individual fitness and population mean fitness, driving evolution by natural selection (Harris et al. 2009; Møller and Mousseau 2015). For example, Moller and Mousseau (2016) reviewed 17 studies that claimed adaptation to radiation in the Chernobyl Exclusion Zone. Four to seven of these studies showed data consistent with an adaptive evolutionary response to radiation but most of the 17 studies suffered from minimal spatial replication, variation in the amount of radiation exposure studied, and were understandably comparative, lacking the careful design of reciprocal transplants or common gardens (Møller and Mousseau 2016). Moller and Mosseau therefore urged new studies that include extensive sampling from multiple sites varying in radiation exposure and they advocated study of multiple generations to rule out epigenetic or maternal effects (Møller and Mousseau 2016).

Goodman et al. (2019) addressed many of Møller and Mosseau’s criticisms in a study of *Daphnia* water fleas, a planktonic crustacean. They collected live *Daphnia* from eight lakes in the Chernobyl Exclusion Zone that varied in radiation exposure. *Daphnia* were then lab-reared under standard, non-radioactive conditions to measure fecundity and survival by third generation female clones. Goodman et al. (2019) found minimal effects of radioactive dose rate on lab-raised *Daphnia* survival, fecundity, and population growth rate. Instead, most of the variation in those fitness proxies was explained by ecological variation from lake to lake (Goodman et al. 2019). This null result either suggests that radiation exposure did not influence evolution; or, that it may be difficult to detect fitness differences in present-day populations if strong, radiation-induced selection caused rapid adaptation that equilibrated differences in mean fitness between populations exposed to variable radiation (Goodman et al. 2019, see also Esnault et al. 2010; Galván et al. 2014). Thus, it may be helpful to test for the effects of radiation in a time series, following the same population as/if it evolves. This temporal approach would control for spatial variation in environment, as well as variable evolutionary history—each lineage would serve as its own control.

Such an approach may be possible through resurrection ecology (Orsini et al. 2013; Weider et al. 2018), wherein a time series of dormant eggs is recovered from annually deposited sediments. If these eggs are hatched and measured for fitness proxies, we may be able to detect the mutation-rate and evolutionary effects of irradiation during and after fallout events. *Daphnia* are useful in this regard (Burge et al. 2018) because their cyclical parthenogenic life history produces annual cohorts of overwintering, diapausing eggs (Kerfoot and Weider 2004). These eggs are stored in a modified carapace, the ephippium, which, if buried during annual sedimentation, can keep eggs viable for decades or even centuries (Burge et al. 2018). Hatching this egg bank reveals a snapshot of population ecology and adaptation at the time of deposition. *Daphnia* are especially useful because their ability to reproduce asexually allows for repeatable, controlled laboratory experiments. Experiments with resurrected clonal lineages have demonstrated adaptation to factors such as novel toxic algae (Hairston et al. 1999), eutrophication (Frisch et al. 2014), temperature change (Geerts et al. 2015), metal pollution (Rogalski 2015, Rogalski 2017), and invasive species (Einum et al. 2022; Rani et al. 2022).

Here, we report a study that aimed to resurrect and measure fitness for *Daphnia* exposed to nuclear fallout from weapons testing. The United States Department of Energy (DOE) made one hundred atmospheric (*i*.*e*., above ground) nuclear weapons tests at the Nevada Test Site (NTS) from 1951 to 1962, before the Partial Test Ban Treaty drove tests underground in 1963 (National Nuclear Security Administration Nevada Field Office 2015). During this nuclear era, fission products showered ~44% of Utah, ~44% of Nevada, and ~54% of Arizona with doses of ^137^Cs and other more active isotopes (Beck 1999; Szymendera 2020). The total estimated activity of ^137^Cs deposited in the contiguous United States from NTS testing is 2.3 petabecquerel (PBq), on the order of radiation released during the nuclear accident at Fukushima (7-20PBq; IAEA 2015) and one order less than that released by Chernobyl (85PBq) (Henri 2002). Thus, downwind areas received a radiation dose that was likely deleterious, certainly at least to the bodies of some human individuals who were dosed. That is, a cohort of people in Utah, Nevada, and Arizona receive compensation if they develop some cancers (Beck 1999; Szymendera 2020).

At least sixty generations of *Daphnia* have passed in the roughly 60 years since aboveground testing ceased. We attempted resurrection ecology to track ~120 years of *Daphnia* life history through time within individual lakes. We predicted that radiation would have caused an increased frequency of mutations deleterious for development, causing survival and reproductive rates to drop during and after testing, relative to rates before testing (Blaylock 1969; Cooley 1973). We expected that survival and reproductive rates would recover over time, as populations repaired damaged DNA, purged deleterious alleles, adapted to compensate for deleterious alleles, or received migrants from less irradiated lakes.

## Methods and Materials

### Preliminary collections and radioisotope dating

We conducted preliminary field work at eight alpine lakes in Utah, USA, in June 2020 (Table 1). We used an inflatable boat and a plankton tow with 64 micron mesh to take at least one vertical plankton tow from each lake to record the presence of life *Daphnia spp*. in each lake. *Daphnia spp*. were obvious in Fish, Lebaron, and Panguitch Lakes, but not Bear, Mirror, Moosehorn, Puffer, or Teapot Lakes. We also used an Ekman dredge to collect samples of the top few cm of each lake bottom. We kept the Ekman samples in the dark, on ice in the field. Several weeks later, in the lab, we isolated ephippia from these samples using dissection microscopy. We placed the ephippia individually in 48-well cell culture plates in Hebert’s Pond Water, illuminated on light boxes at room temperature to promote hatching (Walsh and Post 2011; Walsh and Post 2012) to hatch ephippia. We successfully hatched *Daphnia* ephippia from three of those lakes: Mirror Lake, (Uinta-Wasatch-Cache National Forest), Puffer Lake (Fish Lake National Forest), and Panguitch Lake (Dixie National Forest). Each lake is a natural lake and therefore existed during nuclear testing, though Panguitch and Puffer lakes were dammed. The lakes have not gone dry in their recorded history.

**Table 1.** Coordinates and altitudes Utah lakes visited in 2020. The * indicates lakes with successful *Daphnia* hatching from Ekman grabs. * lakes were radioisotope dated and re-sampled in 2021 for the life history experiment. * lakes were used for stable isotope analysis. * lakes with depths reported for samples taken in 2021, otherwise 2020 values unless not recorded (NR) in the field.

| Lake Name | Latitude | Longitude | Altitude<br>(m) | Depth<br>(ft) |
| --- | --- | --- | --- | --- |
| Bear | 41.9602° | -111.3944° | 1804 | 50 |
| Fish | 38.5388° | -111.7267° | 2696 | 30 |
| Lebaron | 38.2255° | -112.3997° | 3013 | NR |
| Mirror* | 40.7052° | -110.8876° | 3058 | 35 |
| Moosehorn | 40.6957° | -110.8937° | 3167 | 10 |
| Panguitch* | 37.7143° | -112.6407° | 2502 | 34 |
| Puffer* | 38.3165° | -112.3628° | 2947 | 5 |
| Teapot | 40.6805° | -110.9428° | 3031 | 33 |

During preliminary field work, we also collected “gravity” cores with an HTH sediment corer with core tubes 66mm in interior diameter and 40cm long (Renberg and Hansson 2008). Once collected, within six hours in the field, we used the HTH extrusion platform (Renberg and Hansson 2008) and a plexiglass scraper to divide cores into 0.50cm sections, each section placed into individually labeled, 7oz Whirl-Pak sample bags, where, for example, the bag labeled “0.00cm” contained sediment from 0.00-0.50cm. Samples were stored in dark coolers on ice in the field, and at 5C once back at Loyola University Chicago. We used these cores to get radioisotope dating profiles for Mirror, Puffer, and Panguitch Lakes. To accomplish this, we freeze dried and massed sediment samples at the Northwestern University Stable Isotope Biogeochemistry Laboratory and then sent them to St. Croix Watershed Research Station for dating. We used ^210^Pb radioisotope dating on each lake. We also used ^137^Cs radioisotope profiles to date Mirror Lake for comparison to the ^210^Pb dating profile and approximately quantify the amount of radioactive particles that fell on the lakes.

Lead-210 was measured at 15-20 depth intervals in each core through its grand-daughter product ^210^Po, with ^209^Po added as an internal yield tracer following Eakins & Morrison (1978) on an Ortec Octect alpha spectroscopy system (Ortec, USA). Unsupported ^210^Pb was calculated by subtracting supported activity from the total activity measured at each level; supported ^210^Pb was estimated from the asymptotic activity at depth (the mean of the lowermost samples in a core). Dates were estimated using the constant rate of supply model with errors calculated by first-order propagation of counting uncertainty (Appleby, 2001). Cesium-137 activities were measured in Mirror Lake using gamma spectrometry on an Ortec-EGG high-purity, germanium crystal well detector coupled to a Digital Gamma-Ray Spectrometer (Ortec, USA) based on the methods of Ritchie and McHenry (1973).

### Main Collections

To collect ephippia for our life history experiments, we returned to Mirror, Panguitch, and Puffer Lakes from July 19-21, 2021, to collect four to five sediment cores per lake, again using the HTH sediment corer. Again, in the field, within 6 hours, we sectioned cores into individually labeled, 7oz Whirl-Pak sample bags but at 0.25cm intervals this time, where, for example, the bag labeled “0.00cm” contained sediment from 0.00-0.25cm. We reduced the likelihood of contamination between layers by cleaning the scraper between each layer. Bagged samples were then placed in the dark on ice until they were returned and refrigerated at 5C in the dark at Loyola University Chicago on July 24, 2021. We chose two 2021 cores from each lake to radioisotope date and collect ephippia.

We dated each core used for ephippia collection, to account for any within-lake, core-to-core variation. ^210^Pb dating was again conducted by the St. Croix Watershed Research Station following the same methods outlined above. We selected samples every 0.5 cm down to the to the depth where only supported (background) ^210^Pb could be detected in the previous cores and every 1.5 cm below that. Because average wet mass for sediment was ~8g per section, we had sediment left over for isolating ephippia from the same sections that were selected for dating. We again freeze dried the sediments at the Northwestern University Stable Isotope Biogeochemistry Laboratory. We massed them post-drying to calculate percent dry mass and porosity.

### Ephippia Collection

In Fall 2021, while 2021 core sediments were being prepared for radioisotope dating, we collected ephippia from the same cores from each lake. We focused on the above-ground nuclear era (1951-1962) and ~40 years on either side, as estimated from 2020 core dating. For Mirror Lake, ^210^Pb dating of the preliminary 2020 cores suggested that the date range of 1911-2002 fell between 2.50cm-10cm. For Panguitch Lake, depths sampled were 9.50cm-27.00cm. For Puffer Lake, depths sampled were 2.5-10cm.

For each sample in those intervals (*i*.*e*., each Whirl-Pack bag, at 0.25m resolution), sediment was suspended with Millipore-filtered water, then poured in into a 5.5in diameter x 1.0in depth glass petri dish. We examined plates under Leica KL300 dissection microscope at 1 to 4X magnification. Each dish had a paper guide printed with concentric circles and a radius, taped to the bottom, to help ensure that all parts of the plate were assayed. We used forceps to transfer individual ephippia to individual wells of 48-well cell culture plates. Wells were filled with COMBO, a growth medium suitable for *Daphnia* (Kilham et al. 1998). For each ephippium, we noted date examined, lake, core, sample, number of visible eggs, whether the ephippium was floating, and whether there was visible damage to the ephippium or eggs. Between samples, sediment was emptied into waste containers and glass plates were wiped clean. Waste sediment was autoclaved and trashed. Once a well-plate was filled, it was closed with an aluminum plate sealer, the lid was taped on, and the plate was placed in the freezer at -20C.

### Life History Experiments

In late January 2022, ephippia-filled plates were driven on ice over two days from Loyola University Chicago to University of Texas at Arlington. On January 25^th^, 2022, frozen plates were unsealed and placed in an incubator at 16C and 14:10 light-dark photoperiod. All ephippia were evaluated daily for hatching (following Walsh and Post 2011; Walsh and Post 2012). Because of evaporation, we added COMBO to wells as needed. Nothing had hatched by February 4, 2022, so we resealsed and refroze the eggs. On February 7, we thawed plates and placed them back on the incubator. Only a single ephippium had hatched by 21 February, so we refroze the plates. On February 27, we placed the plates back in the incubator and changed the photoperiod to 24-hour light. We monitored until March 1, 2022, but nothing hatched. Chlorine treatment has been shown to improve hatching success (Paes et al. 2016). As a trial, we chlorine treated 10% of the ephippia: for six wells per plate, we soaked ephippia in a 20% chlorine solution for three minutes, rinsed the ephippia with DI water, soaked them in DI water for three minutes, and then returned them to their wells in COMBO. We monitored all ephippia until March 11, 2022. Because there was still no hatching, we halted the experiment, resealed and refroze the plates and drove them back to Loyola University Chicago for storage at -20C, where the plates remain.

### Stable Isotope Analysis

At the same time as United Stats above ground nuclear testing, alpine lakes started being affected by expanding mining, industry, and agriculture, changing stoichiometric and chemical conditions in freshwater systems (Vitousek et al. 1997). Thus, water quality and selection regimes for *Daphnia* communities in our lakes may have changed concurrently with nuclear testing (Wolfe et al. 2001; Wolfe et al. 2003; Reynolds et al. 2010), confounding our efforts to understand how irradiation influences evolution. To assess this potential confounding variable, we also prepared ephippia and sediments from 2020 cores for stable isotope (δ^15^N and δ^13^C) analysis.

For one core from each of Mirror, Panguitch, and Puffer Lakes, we sampled sediment sections that spanned the core. For each sample, we collected as many ephippia as possible and pooled them into a single, labeled PCR tube. The pooled wet mass was recorded for each section. Then ephippia in those tubes were freeze dried, massed, and packed in tin capsules for analysis on a Costech 2010 Elemental Analyzer coupled to a Delta V Plus isotope ratio mass spectrometers (EA-irmMS) at the Stable Isotope Biogeochemistry Laboratory at Northwestern University. To estimate background change in total organic carbon (TOC) and total organic nitrogen (TOC) stable isotope ratios, we followed the same protocols (with an added acidification step to remove inorganic carbon from our sediments) to prepare sediment samples spanning each core for analysis at the Stable Isotope Laboratory. Then we visualized change through time in *Daphnia* stable isotope ratios, relative to sediment background total organic carbon and nitrogen.

## Results

### Core Chronology

^210^Pb chronologies are presented in Figure 1 for one core from each lake in 2020 and two cores from each lake in 2021. Gravity cores reached the background detection limit in all three lakes. The ^137^Cs profile from the 2020 Mirror Lake core showed two peaks (Figure 2). The later, larger peak corresponds to the 1963 global ^137^Cs maximum. The earlier peak dates to the early 1950s and may represent local fallout from earlier testing. The average of the mean sedimentation rate across three cores for Mirror Lake was 0.012 g/cm^2^/year (s.d. of means = 0.006). The average of the mean sedimentation rate across three cores for Panguitch Lake was 0.031 g/cm^2^/year (s.d. of means = 0.006). The average of the mean sedimentation rate across three cores for Puffer Lake was 0.027 g/cm^2^/year (s.d. of means = 0.002). The raw data from radioisotope dating are provided on Data Dryad (https://doi.org/10.5061/dryad.83bk3j9tx).

**Figure 1.**
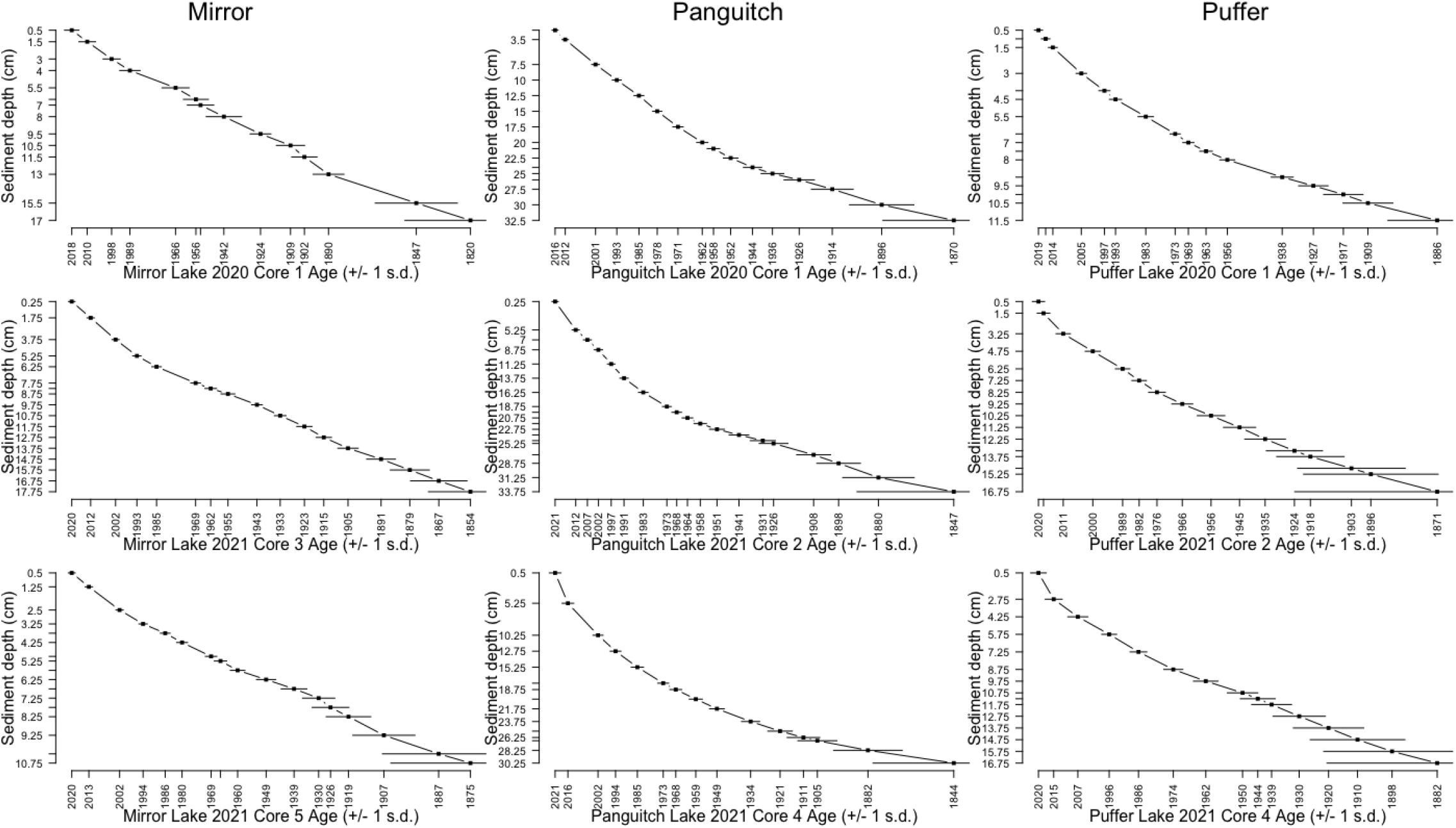
Depth (cm) versus year (AD) profiles for Mirror (column 1), Panguitch (column 2) and Puffer (column 3) lakes, based on ^210^Pb radioisotope dating.

**Figure 2.**
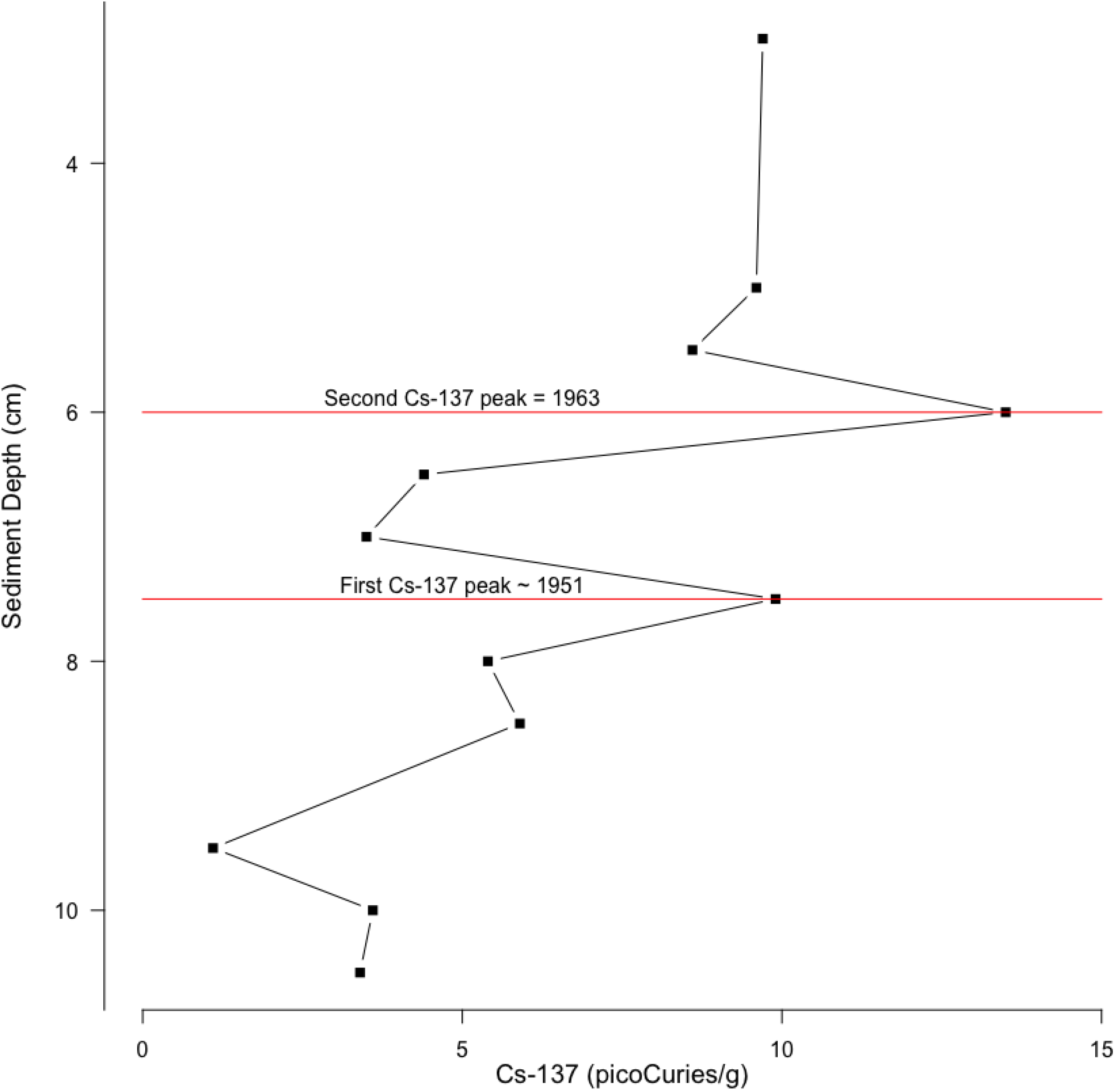
^137^Cs levels against depth in Mirror Lake, from a single core sampled in 2020. A value of 10 picocuries translates to approximately 0.37 radioactive decays per gram of sediment per second.

### Ephippia

We isolated 4716 ephippia in total across 6 cores, 2 cores from each lake, taken in 2021. Table 2 shows ephippia numbers by lake and core. Figure 3 shows number of ephippia recovered against inferred time.

**Table 2.** Ephippia counts by lake and core, from 2021 sampling.

| Lake Name | Core ID | Number of<br>ehippia |
| --- | --- | --- |
| Mirror | 03 | 1290 |
| Mirror | 05 | 396 |
| Panguitch | 02 | 1113 |
| Panguitch | 04 | 965 |
| Puffer | 02 | 846 |
| Puffer | 04 | 106 |

**Figure 3.**
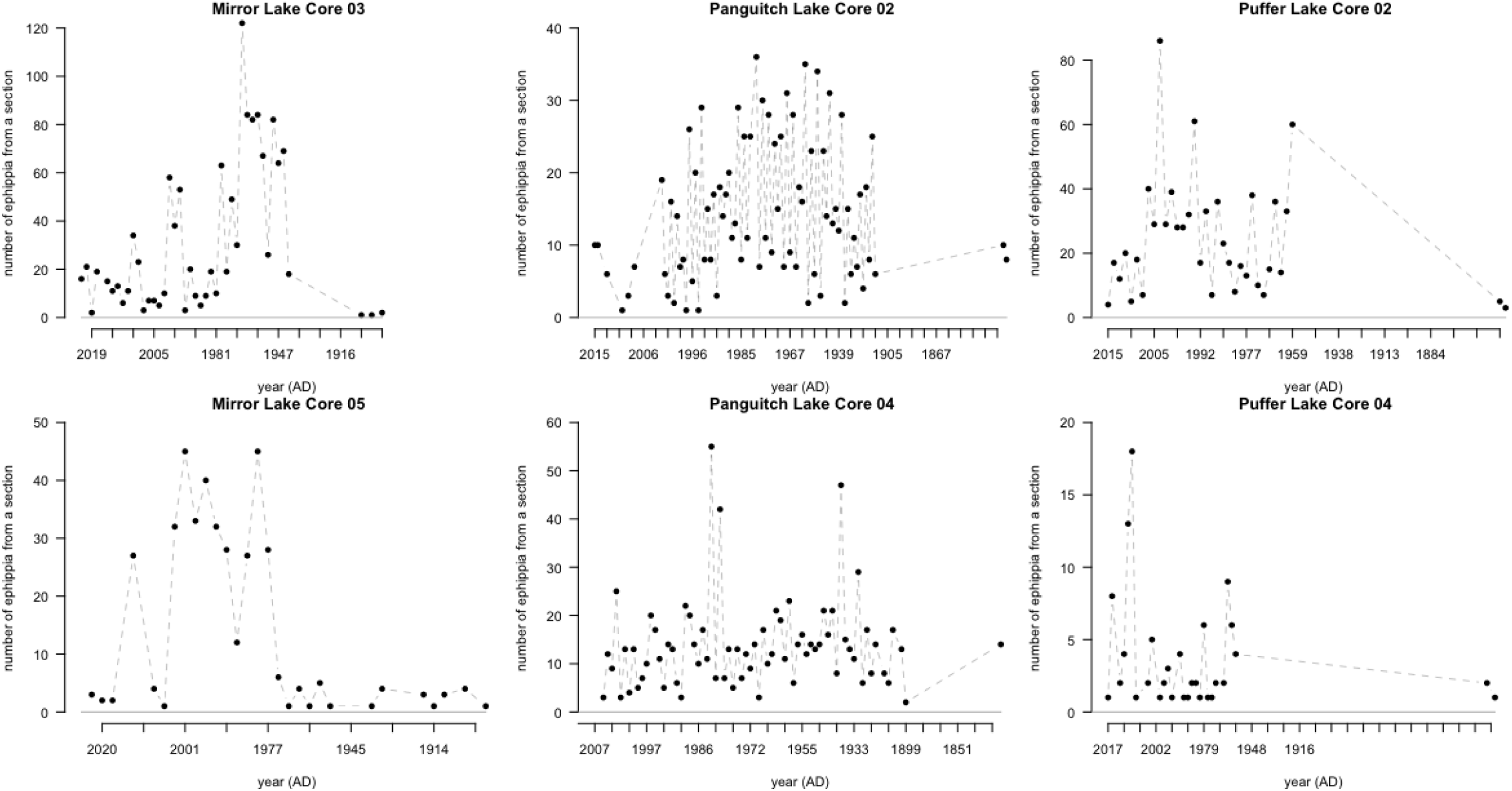
Ephippia recovered by section per core. We sampled sections, at 0.25cm resolution. Points represent counts for samples where we looked for and found ephippia.

### Life history experiment

Of the 4716 ephippia, only one hatched, from Puffer core 02, 9.50cm. Ironically, radioisotope dating infers that 9.50cm corresponds to ~1960, while above ground nuclear testing at the Nevada Test Site was still active. That individual produced 3 clutches in 7 days, with respective clutch sizes of five, nine, and twelve offspring, 26 total. This suggests that it if these eggs could be hatched, they would be useful for life history experiments.

### Ephippia and sediment stable isotope change through time

Stable isotope ratios for nitrogen and carbon show appreciable variation through time, though with some trends within and across lakes (Figure 4). Ephippia δ^15^N values decrease in Puffer and Mirror Lakes but stay roughly stable in Panguitch Lake. Ephippia δ^13^C values fluctuate with only an increasing trend in Puffer Lake. Sediment δ^15^N values decrease through time, while Panguitch sediment δ^15^N values stay roughly stable. Sediment δ^13^C values drop in Panguitch lake, while staying roughly stable in Puffer. We cannot make a formal statistical comparison of trends in sediment and ephippia δ^15^N and δ13C values because the samples did not come from the same sections. However, trend lines for Puffer Lake for both δ^15^N and δ^13^C appear positively correlated. For Panguitch Lake, trend lines seem uncorrelated for δ^15^N and possibly negatively correlated for δ13C. We did not measure sediment stable isotopes for Mirror Lake.

**Figure 4.**
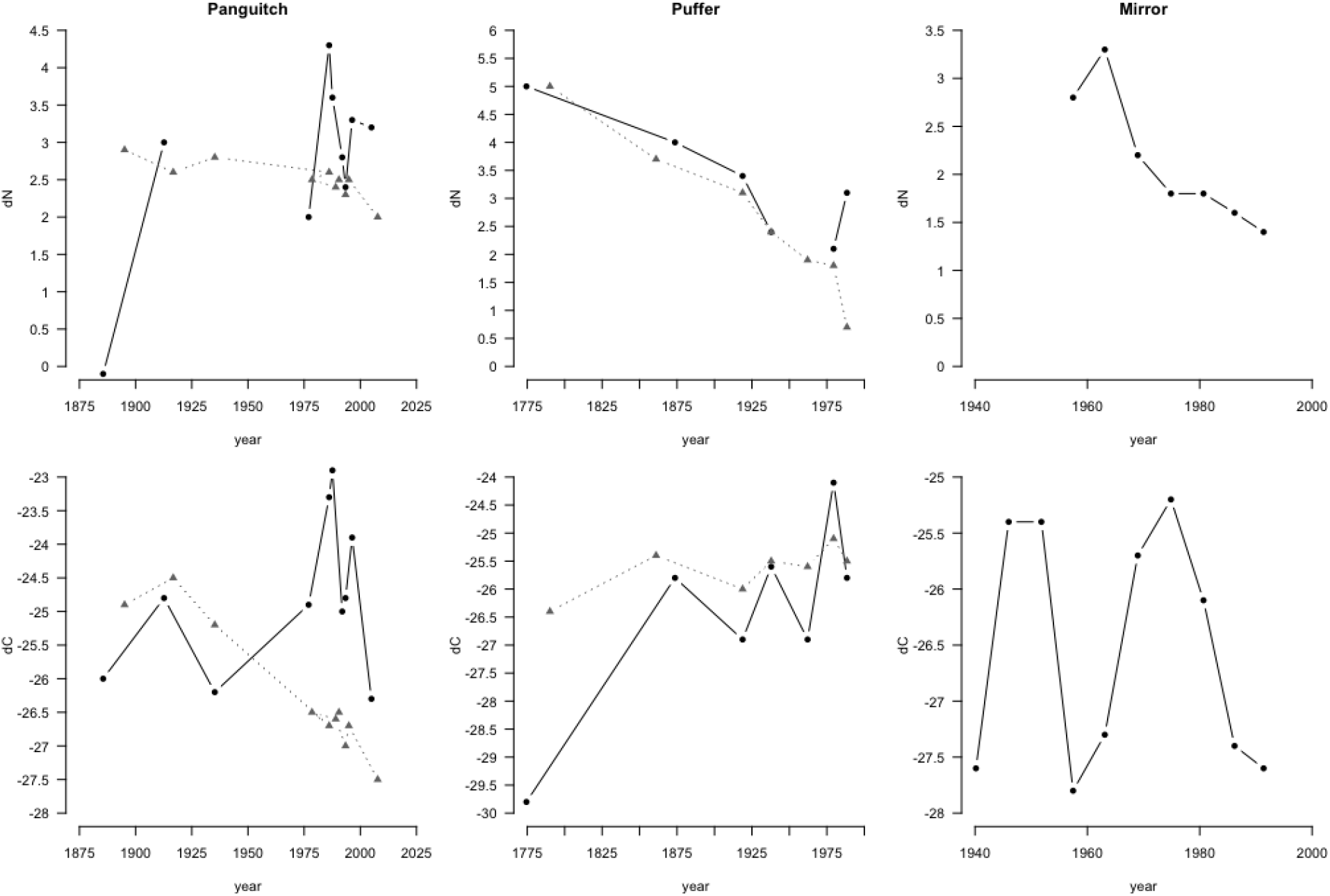
Stable isotope change through time for Panguitch, Puffer, and Mirror Lakes, in δ^15^N ratios (top row) and δ^13^C ratios (bottom row). Black circles denote values for ephippia, whereas gray triangles represent bulk sediments. Bulk sediment data were not collected for Mirror Lake. Gaps in ephippia δ^15^N values represent samples that were not successfully analyzed.

## Discussion

We took sediment core samples from three alpine, Utah lakes that experienced ionizing radiation during aboveground nuclear weapons testing at the Nevada Test Site from 1951-1962. ^210^Pb radioisotope dating showed that those cores dated back to the 1800s, giving us access to an egg bank that spanned the nuclear era and making possible a resurrection ecology experiment to test the effects of nuclear fallout on evolution. We isolated more than 4700 ephippia from those cores and used standard and new techniques to try to hatch the eggs. Unfortunately, we only managed to hatch one—deposited during the nuclear era. Thus, our experiment failed, and we could not answer our research questions.

Why didn’t the ephippia hatch? We can speculate several reasons. These high-altitude lakes might experience slower sedimentation rates than low-altitude lakes, resulting in a smaller likelihood that ephippia are covered by sediment. As such, there may be higher hatching rates each spring, leaving fewer ephippia dormant and available to resurrection experiments.

Moreover, high-altitude lakes likely experience larger extremes in environmental conditions, especially when shallow like Puffer and Mirror Lakes, with frequent mixing and more freezing, and larger changes in water levels. In 2020, we sampled cores from Puffer at ~24ft deep; the same spot in 2021 was only 5ft deep. These factors may cause faster degradation of dormant ephippia. Such speculation does not explain resurrection failure in Panguitch Lake, which was lower altitude, much larger, dammed, and had a robust, modern population of *Daphnia pulicaria*, based on plankton tows. Personal communications from researchers in this field suggests that nearly everyone has had a resurrection study fail because hatching failed, for whatever reason.

Thus, we present this manuscript because null results should be published, and because this is a venue to make our radioisotope dating and sedimentation rate data available to paleolimnologists studying environmental change in lake systems through time (https://doi.org/10.5061/dryad.83bk3j9tx). We also present preliminary stable isotope data for *Daphnia* ephippia and lake-bottom sediments, which showed appreciable variation through time.

## Acknowledgements

We thank C.J. Thompson of the University of Texas at Austin and Los Alamos National Labs for help with study conception and grant writing. We thank C. Fix, F. Joaquin, M. Marino, S. Swank, M. Walsh, M. Korte, A. Christian, and C. Miller for help with field work, sample preparation, data collection, and the life history experiment. We thank E. Mortenson of the St. Croix Watershed Research Station for radioisotope dating. We thank R. O’Grady for advice and help with core collections. We thank the Northwestern University Stable Isotope Biogeochemistry Laborabory and Y. Axford of the Northwestern University Department of Earth and Planetary Sciences for help with study design, sediment preparation, and stable isotope analysis. Permission to collect sediment cores was granted by United States Forest Service rangers of the Dixie, Fish, and Uintas-Wabash-Cache National Forests. This work was funded by a National Science Foundation EAGER award (DEB-2028775) to Y.E.S and M.R.W.

